# Investigation of cell metabolism dysfunction in autism spectrum disorders using genome-scale metabolic network analysis of neuronal development

**DOI:** 10.1101/2024.10.01.616152

**Authors:** Moohebat Pourmajidian, Sayed-Amir Marashi

**Author notes:** **Corresponding Author:** Sayed-Amir Marashi.

## Abstract

Autism Spectrum Disorders (ASDs) are a group of highly heterogeneous neurodevelopmental conditions characterized by impairments in social interactions and repetitive behaviors. In spite of extensive efforts, known genetic mutations can hardly explain the ASD cases, which signifies the highly heterogeneous nature of ASDs and calls for shifting to a more holistic approach for understanding the mechanism of these disorders. Several studies have reported that a wide range of metabolic markers is present in blood, urine and brain samples of ASD patients. In this study, using transcriptome data of neurons derived from ASD and control individuals, we reconstructed neuron-specific metabolic network models for two neurodevelopmental time points representing late neural progenitor stage and mature neurons. We simulate gene inactivation in these networks to investigate the altered metabolic fluxes and biological pathways in ASD metabolism. We further examine these networks using reporter metabolite analysis to identify those metabolites around which significant gene expression changes occur. Briefly, our model was successful in predicting previously reported metabolites and pathways implicated in ASD metabolism, including oxidative phosphorylation and fatty acid oxidation, and points to a new finding concerning alterations in the less studied pathways like keratan sulfate degradation pathway and N/O-glycosylation in ASD. Finally, by employing the robust metabolic transformation algorithm (rMTA) we set to find potential drug targets that can reverse the ASD metabolic state into a healthy one. Interestingly, our model found phosphatidic phosphatase enzyme, as well as phosphatidylethaolamine and phosphatidylserine transfer proteins as targets that could recover the healthy metabolic state.

## Introduction

### Autism Spectrum Disorders

Autism Spectrum Disorders (ASDs) are a group of highly complex neurodevelopmental disorders characterized by deficits in social interactions and repetitive behaviors (Geschwind, 2011; De La Torre-Ubieta et al., 2016; Ansel et al., 2017; Vorstman et al., 2017). There is a significant genetic and phenotypic heterogeneity among affected individuals, which manifests in a broad range of symptom severity. The prevalence of reported ASD cases has been rising drastically and it is estimated to affect around %1 of the population globally (Lord et al., 2020).

With the revolutionary advances in genomic techniques and methodologies such as whole-exome sequencing and single-cell RNA-sequencing, extensive research has been done to decipher the genetic mutations and neurobiological mechanisms underlying ASD. Several studies report that many of the genes implicated in ASD converge into a set of specific biological pathways (Voineagu et al., 2011), including synaptogenesis, protein synthesis and degradation, and chromatin remodeling (Pinto et al., 2014; Bourgeron, 2015). Despite a considerable amount of research and large cohort studies, known genetic mutations are estimated to collectively account for only about 10%-20% of the cases, with none of them individually accounting for more than 1% of ASD cases (Jeste and Geschwind, 2014). This, in turn, signifies the highly heterogeneous nature of ASD and calls for shifting to a more holistic approach for studying these disorders.

### ASD and metabolic dysfunctions

Metabolic dysfunction in ASD patients was first reported in the 80’s in terms of increased lactate levels in the plasma (Coleman and Blass, 1985). Later in 1998, Lombard proposed that ASD may be “a disorder of atypical mitochondrial function” after observing several metabolic abnormalities in ASD patients including lactic acidosis and decreased level of ATP in the brain (Lombard, 1998). Mitochondria, aside from supplying the ATP demand of the body, play a critical role in calcium homeostasis and signaling, production of reactive oxygen species, and regulation of cell apoptosis. Mitochondria have been implicated in several neurodevelopmental processes including neuronal differentiation and maturation, formation of dendritic processes, cortical migration, and synaptic plasticity (Hollis et al., 2017). The human brain accounts for approximately 20% of the body’s energy demand and neurons rely on great amounts of ATP for maintaining ionic gradients for neurotransmission and synaptogenesis (Cheng et al., 2017). Over the last 40 years, several research groups have been investigating metabolic and bioenergy dysfunction in ASD. The prevalence of mitochondrial diseases is estimated to be 5% in ASD patients compared with 0.01% in the general population (Rossignol and Frye, 2012; Cheng et al., 2017). Multiple reports show the presence of abnormal metabolic biomarkers and oxidative phosphorylation defects in blood, urine, skeletal muscle, and brain of ASD individuals (Rossignol and Frye, 2012; Tang et al., 2013; Siddiqui et al., 2016; Hollis et al., 2017; Orozco et al., 2019; Žigman et al., 2021). Mutations in genes associated with mitochondrial function like NDUFA5, SLC25A12, and ATP5 family are frequently reported in ASD patients (Chalkia et al., 2017). Therefore, there is a strong body of evidence suggesting that ASD is associated with cell metabolism dysfunction.

### Computational approaches to metabolic dysfunction analysis

With the accumulation of ‘omics’ data on ASD, a systematic investigation of altered metabolic pathways can shed light on the molecular causes of these disorders. Genome-scale metabolic network models (GEMs), owing to biological and biochemical knowledge, can provide a suitable platform for studying implicated metabolites and pathways in health and disease (Cook and Nielsen, 2017; Fouladiha and Marashi, 2017; Marashi, 2017; Sertbas and Ulgen, 2018). GEMs can link gene-protein-reaction knowledge to the activity of metabolic pathways. In other words, the application of cell-specific metabolic network models enables us to understand how the “reductionistic details” of genetics can affect its “holistic” metabolic phenotypes.

In the present study, using publicly available transcriptomic data, we first reconstruct genome-scale metabolic models of human iPSC-derived neurons in two distinct developmental stages, *i.e.*, late neural progenitor (Day 35) and mature neurons (Day 135). In the next step, we determine those genes that are differentially expressed between ASD and control individuals, and carry out *in silico* knockout at each developmental stage to reconstruct metabolic models representing ASD metabolism. Then, we investigate the resulting metabolic alterations to understand how gene regulation affects the active metabolic pathways in ASD vs. normal cells.

## Materials and Methods

### Data acquisition and identification of differentially expressed genes (DEGs)

The RNA-seq data used in this study was reported in DeRosa et al. (2018) (DeRosa et al., 2018). This dataset contains single-cell RNA-Seq data of 6 male individuals with idiopathic ASD and 5 male control individuals. RNA sequencing has been performed on iPSC-derived neurons at two time points after “cortical neurogenesis” induction: 35 days, representing late neural progenitor (LNP) stage; and 135 days, representing mature neuron (MN) stage. Raw HTSeq counts were downloaded from Gene Expression Omnibus (GEO ID: GSE124308). Normalization and differential expression analysis were performed using DESeq2 package in R (Love et al., 2014). We selected differentially expressed genes (DEGs) with adjusted *p*-value<0.05 (after Benjamini-Hochberg correction), based on the fold change threshold of Log_2_(*FC*)<-0.8.

### Model reconstruction

Figure 1 schematically represents the reconstruction and analysis of the cell-specific networks used in this study. The generic human metabolic reconstruction Recon 2.2 was used in this study (Swainston et al (Swainston et al., 2016)). To reconstruct the neuron-specific metabolic model, transcriptome data were leveraged by employing the CORDA2 algorithm (Schultz et al., 2017). DESeq2 normalized expression values of control subjects were averaged and mapped to Recon 2.2 reactions using mapExpressionToReactions function in COBRA toolbox 3.1 (Heirendt et al., 2019). The CORDA algorithm pipeline requires three groups of reactions as its input: High confidence (*i.e.*, reactions with strong expressional evidence); Medium confidence (reactions with medium-to-low expression levels); and negative confidence (reactions whose activities are not supported by transcriptome data) (Schultz and Qutub, 2016). The mapped expression values were normalized by subtracting the mean and dividing by the standard deviation. Values greater than 5 were considered as high confidence, values between 0 and 5 were considered as medium confidence, while values below zero were considered as negative confidence. When available, Boundaries of the exchange reactions were set according to the culture media composition, Lewis et al. and Çakir et al. (Çakιr et al., 2007; Lewis et al., 2010).

**Figure 1.**
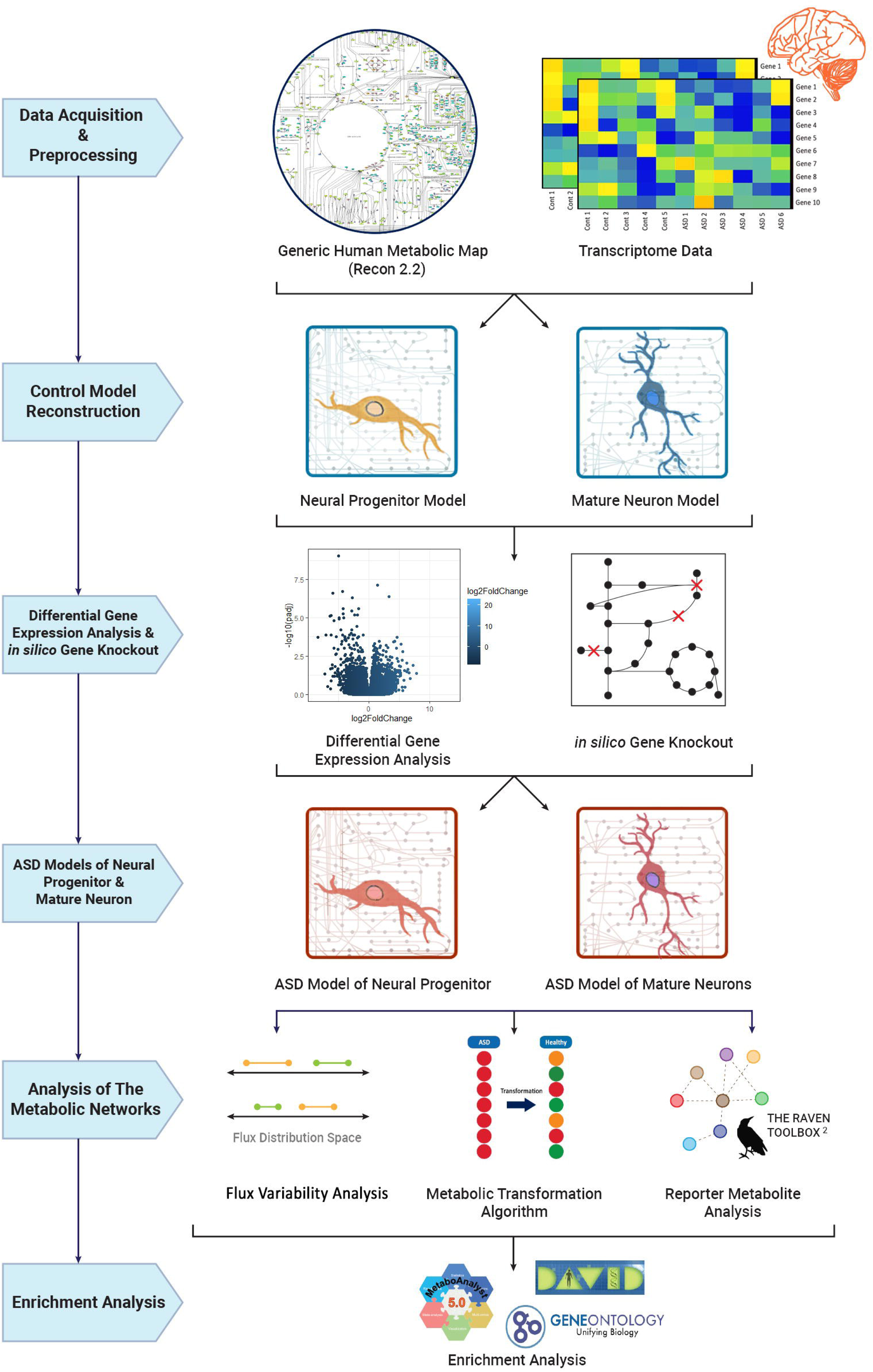
The workflow of the metabolic network model reconstruction and analysis in the current study.

### Flux Balance Analysis

Flux balance analysis is a mathematical approach for calculating flux distributions of the reactions in a metabolic network model. Given a stoichiometric matrix **S** of size *m* × *n* (where *m* represents the number of metabolites and *n* represents the number of reactions in the network), FBA solves a linear program to predict the flux through all reactions in the model represented by the flux distribution vector **v**. By assuming a steady-state condition and other capacity constraints (reversibility and upper and lower flux boundaries), FBA optimizes a given biologically relevant objective function (Orth et al., 2010).

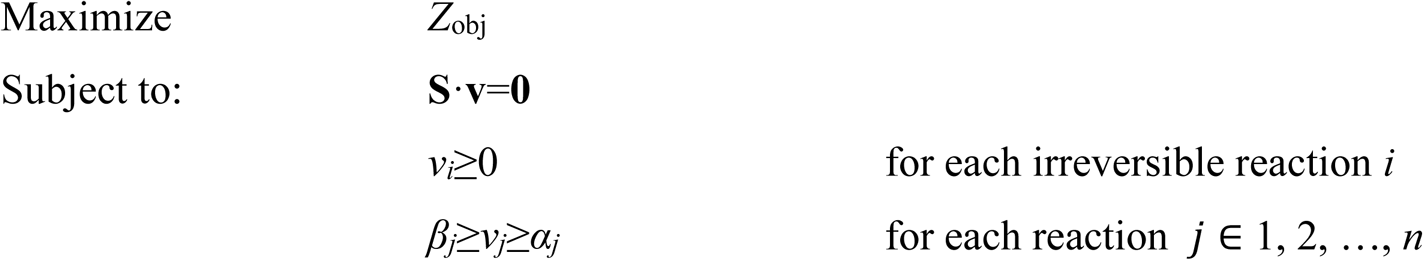

Considering the significant energy demand of neurons and the essentiality of ATP production for neuronal function, maximization of ATP hydrolysis was considered as the objective function of our reconstructed model, that is, *Z*_obj_ = *v*_ATP_hydrolysis_, which ensures maximum ATP production. The solver ‘glpk’ was used for FBA optimization of our models.

### Simulating neuronal metabolism in normal and ASD states

In this study, we focused on understanding the link between gene dysfunction and ASD-related metabolic phenotypes. To this end, we used the above-mentioned “control” metabolic models to obtain ASD models. For each developmental stage (LNP and MN), downregulated DEGs were deleted in their corresponding control models using deleteModelGenes function in the COBRA toolbox to obtain models representing the ASD state. Given a list of genes, this function deletes those genes from the output model and constrains the boundaries of the affected reactions to zero.

Flux variability analysis (FVA) is a method for computing the minimum and maximum flux values of all the reactions in the model while maintaining the optimal value of the objective function (Mahadevan and Schilling, 2003). We performed FVA by employing the fluxVariability function in the COBRA toolbox. To avoid computationally introduced minuscule flux differences, all the FVA outputs were rounded off to 3 decimals to find a flux range for every reaction in each model.

The flux distributions were compared between ASD and control models for both developmental stages. Then, in each model, we determined any reaction whose flux range altered to a new non-overlapping flux range (Yousefi et al., 2019). Genes and subsystems associated with the altered fluxes were extracted. Gene ontology (GO) enrichment analysis was performed using DAVID functional annotation tool v6.8 (Huang et al., 2009a, 2009b).

### Reporter metabolite analysis

We used the reporterMetabolites function in the RAVEN toolbox (Wang et al., 2018) to perform reporter metabolite analysis (Patil and Nielsen, 2005). Differentially expressed genes and their corresponding adjusted *p*-values were used as inputs. Top 20 reporter metabolites with their associated reactions, genes, and subsystems were selected for further analysis. Currency metabolites H^+^, H_2_O were excluded since they mapped to too many reactions in the metabolic model. Metabolite set enrichment analysis (MSEA) was performed using the MetaboAnalyst 5.0 web platform (Xia and Wishart, 2010) when the universal metabolite IDs were available. The reporter metabolites were illustrated using the Escher web application (King et al., 2015).

### Metabolic Transformation Algorithm

To perform the metabolic transformation analysis, we used a recent implementation of the algorithm, called rMTA, in the COBRA Toolbox. This algorithm requires the differential gene expression profile between the source state (*i.e.*, ASD) and the target state (*i.e.*, control) and a reference flux distribution (**v**^ref^) of the source state. Using the ACHRSampler implemented in the COBRA toolbox, 2000 sample fluxes were obtained from each ASD model and the mean was defined as **v**^ref^. Differential gene expressions were given to rMTA along with reference flux of the source state to obtain those perturbations that can shift the metabolic state of ASD towards the target healthy state at the reaction level. Reactions and their corresponding robust transformation score (rTS) were obtained for further investigation.

## Results

### Metabolic network model reconstruction of neural progenitors and mature neurons

In order to reconstruct cell-specific metabolic models of neurons, for each developmental stage, we applied the CORDA2 algorithm. The generic human metabolic network model (Recon 2.2) was pruned using the normalized gene expression data to reconstruct the neuron-specific models. These models are available in Supplementary files S1 and S2. The detailed properties of the models are shown in Table 1. Uptake and secretion rates were constrained to simulate the metabolic environment of the cells. Details of lower and upper bounds used in this study are presented in Supplementary Data Sheet S1. Due to the importance of ATP production in neuronal cells, we added an ATP hydrolysis reaction as the objective function of the models, which amounts to the maximization of ATP production. Then, we performed FBA to ensure that the models were feasible and able to produce ATP.

**Table 1.** Detailed properties of the reconstructed control models. LNP, late neural progenitor stage (day 35); MN, mature neuron stage (day 135).

| Models | Reactions | Metabolites | Genes |
| --- | --- | --- | --- |
| Control LNP | 2093 | 1568 | 927 |
| Control MN | 2231 | 1639 | 957 |

### Differentially expressed gene analysis, FVA, and enrichment analysis

To investigate the metabolic alterations in ASD compared to LNP and MN controls, we carried out flux variability analysis (FVA) to get the flux range of the reactions in each control model and their respective ASD models. Based on our differential gene expression analysis, we obtained a list of all the significantly down-regulated genes in the ASD samples compared to controls (*p*-value < 0.05). Results of the differential expression analysis are available in Supplementary Data Sheet S2. To build a metabolic model for ASD, those genes with a log_2_*FC* < −0.8, were identified and then knocked out in their respective control models. We carried out FVA to compare flux distributions for every reaction between ASD and control models. A Venn diagram (Heberle, Meirelles, da Silva, Telles, & Minghim, 2015) showing the number of shared and unique reactions between the control and ASD models is shown in Figure 2. Figure 3 shows possible outcomes of flux distribution change observed in ASD models compared with control. In this study, those reactions with fully decreased or increased fluxes (condition A and B) were chosen for further analysis and genes associated with these reactions were used for GO biological processes enrichment analysis with DAVID functional annotation tool. A list of non-overlapping altered fluxes (fully decreased and increased) and their associated subsystems at both the neural progenitor and mature neuron stages in our ASD models is provided in the Supplementary Data Sheet S5.

**Figure 2.**
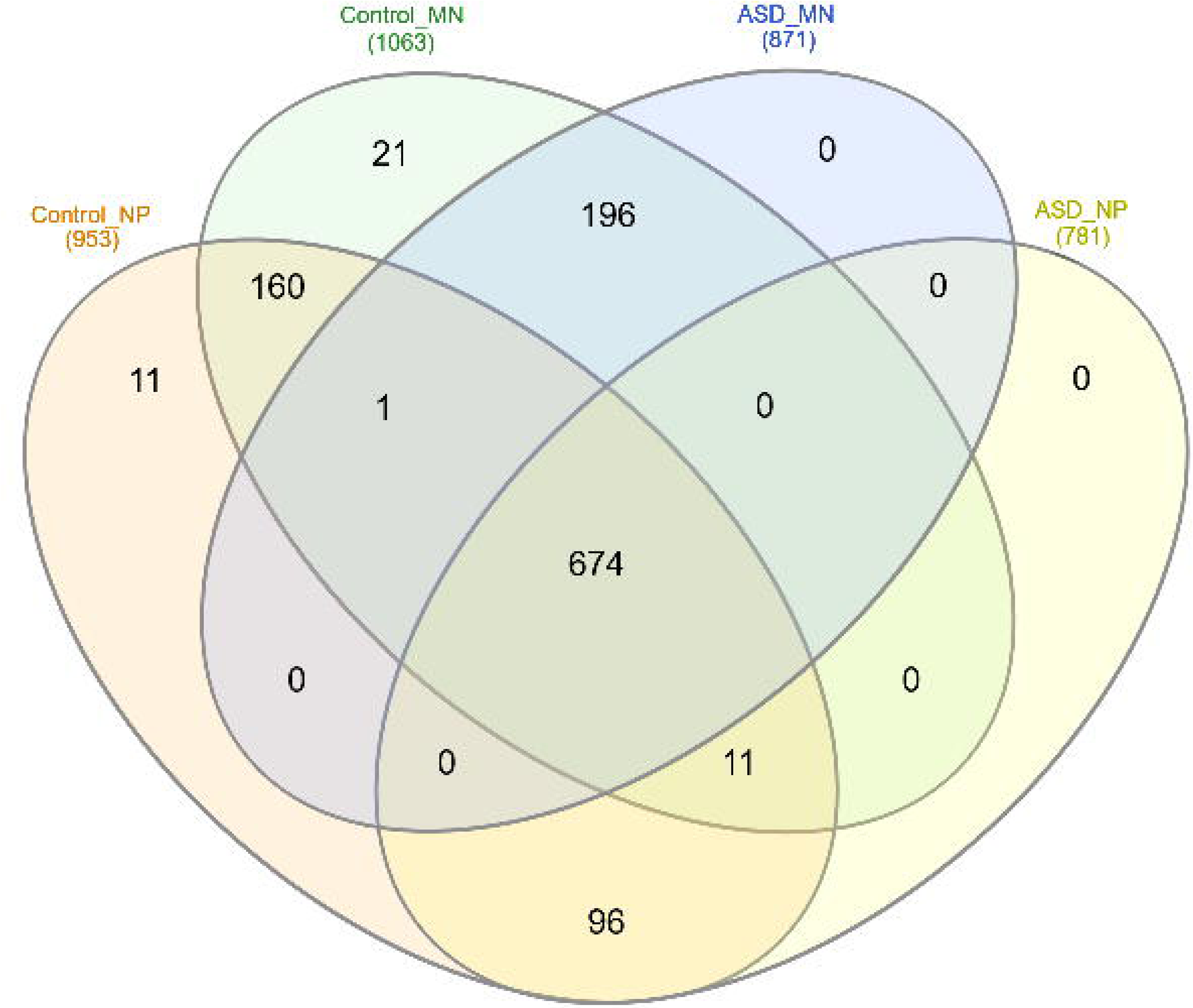
Venn diagram showing the number of shared and unique unblocked reactions between the control and ASD models reconstructed in this study.

**Figure 3.**
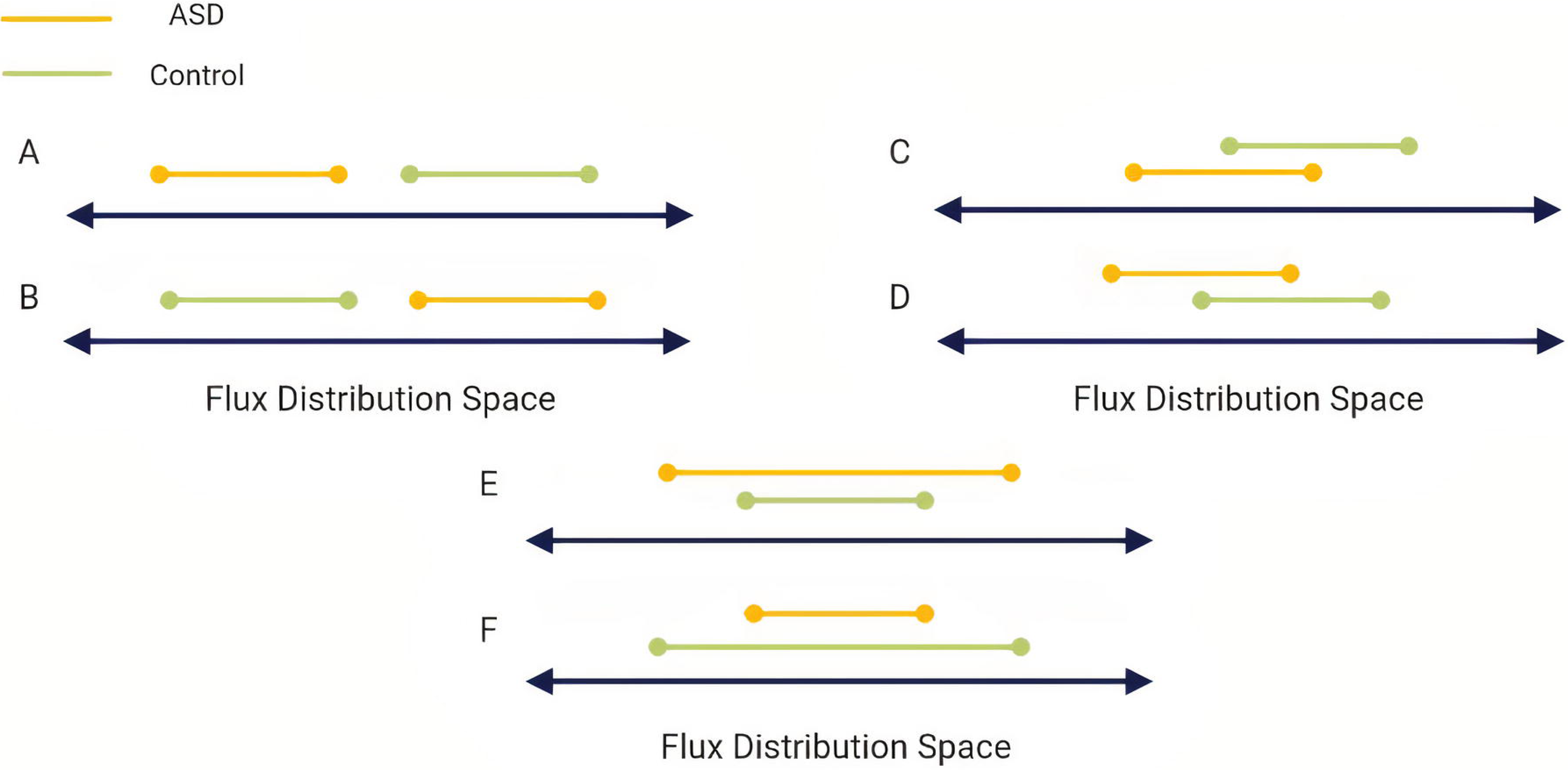
Possible outcomes of flux distribution changes between control and ASD models following FVA analysis. In this study we focused on condition A and B for further investigation.

GO term analysis of the FVA results showed that down-regulated reactions in the ASD models were significantly enriched in oxidative phosphorylation, mitochondrial electron transport, ATP synthesis, and protein N-glycosylation (Fig. 4A). Enrichment analysis also showed that upregulated reactions in ASD significantly correspond to keratan sulfate (KS) metabolism, protein O-glycosylation, transferrin transport, macroautophagy, and phagosome acidification and maturation (Fig. 4B)

**Figure 4.**
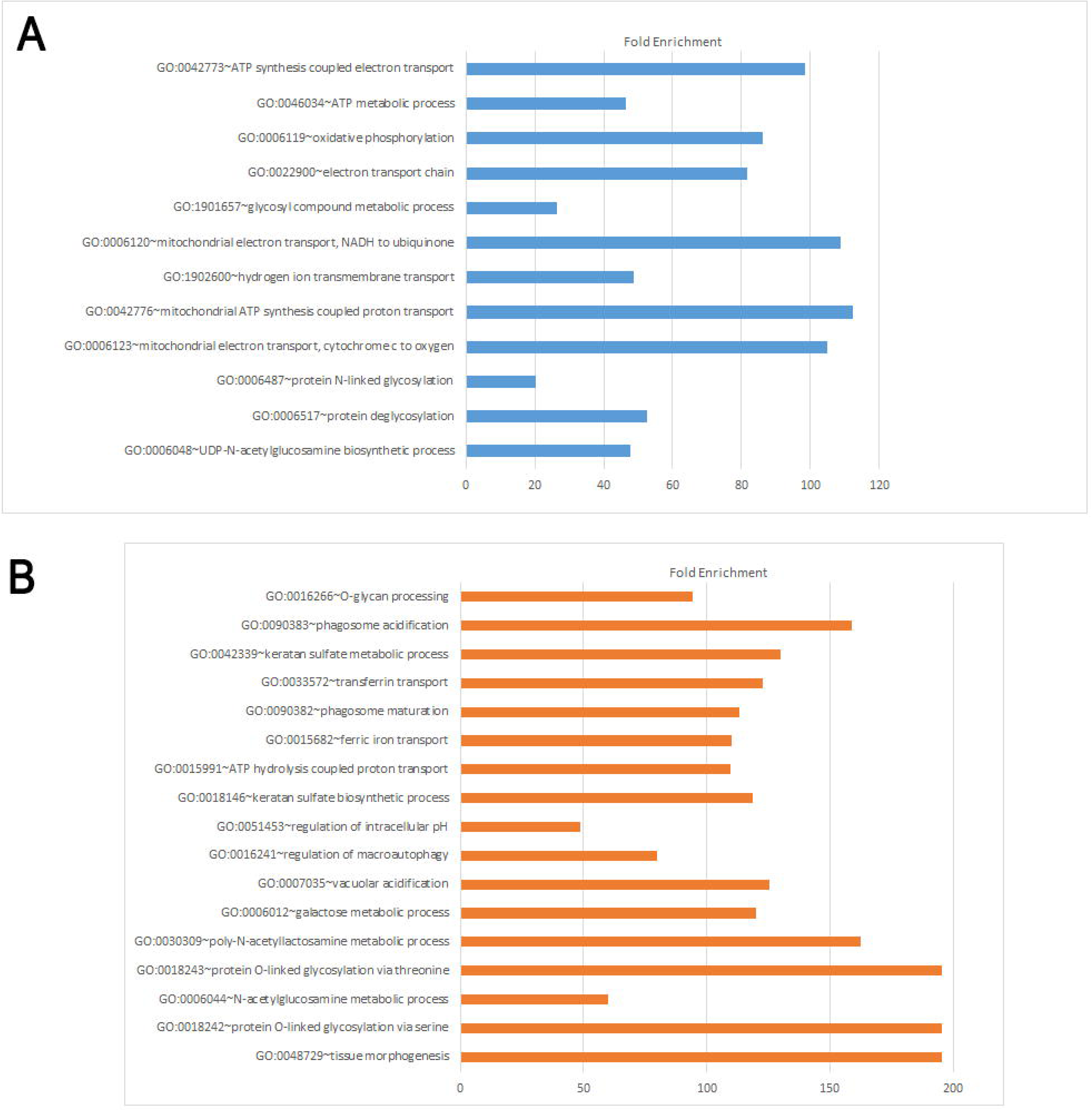
GO biological processes enrichment analysis. GO term enrichment was carried out on FVA results using DAVID functional annotation tool. (A) Down-regulated reactions in ASD models; (B) Up-regulated reactions in ASD models. The order indicates significance based on FDR values.

### Reporter metabolite analysis

We performed reporter metabolite analysis to find significantly affected metabolites in the network based on differential gene expression analysis between ASD and control samples. By incorporating the control models, their respective DEGs and *p*-values, a list of reporter metabolites was obtained using Raven Toolbox. Analysis was done considering three cases: all the DEGs, only up-regulated DEGs, and only down-regulated DEGs. Typically, the top 20 metabolites are considered reporter metabolites (Table 2). A full list of reporter metabolites is presented in the Supplementary Data Sheets S3 and S4.

**Table 2.** Top 20 reporter metabolites obtained by reporter metabolite analysis at LNP and MN developmental stages.

| Neural progenitor model | Mature neuron model |
| --- | --- |
| 3,5-cyclic AMP (cAMP) | D-galactose |
| trans-2-cis-8-tetradecadienoyl-CoA | N-acyl-beta-D-galactosylsphingosine |
| 3,6,9-dodecatrienoylcoa | 5,6-dihydrothymine |
| 3,6,9,12-octadecatetraenoylcoa | thymine |
| (3Z,6Z)-dodecadienoyl-CoA(4-) | F1alpha |
| 2,7,10,13,16-docosapentenoylcoa | keratan sulfate I, degradation product 11 |
| 2,7,10,13-hexadecatetraenoylcoa | keratan sulfate I, degradation product 14 |
| 2,7,10-hexadecatrienoylcoa | keratan sulfate I, degradation product 17 |
| 2,6,9,12-octadecatetraenoylcoa | keratan sulfate I, degradation product 20 |
| coenzyme Q10 | keratan sulfate I, degradation product 23 |
| (3S)-3-hydroxylinoleoyl-CoA | keratan sulfate I, degradation product 26 |
| (3S)-3-hydroxy-cis-8-tetradecenoyl-CoA | keratan sulfate I, degradation product 29 |
| ubiquinol-10 | keratan sulfate I, degradation product 32 |
| (2S,3S)-3-hydroxy-2-methylbutanoyl-CoA(4-) | keratan sulfate I, degradation product 35 |
| 3(S)-hydroxy-5Z,8Z-tetradecadienoyl-CoA | keratan sulfate I, degradation product 38 |
| (3S)-3-hydroxy-cis,cis-palmito-7,10-dienoyl-CoA | keratan sulfate I, degradation product 39 |
| (3S)-3-hydroxydodec-cis-6-enoyl-CoA | keratan sulfate I, degradation product 40 |
| 3-hydroxy-2-methylpropanoyl-CoA | keratan sulfate I, degradation product 41 |
| trans-2-cis,cis-5,8-tetradecatrienoyl-CoA | keratan sulfate I, degradation product 8 |
| deacylated-(phosphoethanolaminyldimannosyl),(phosphoethanolaminyl)-mannosylglucosaminyl-acylphosphatidylinositol | keratan sulfate II (core 2-linked), degradation product 5 |

Our analysis showed that at the LNP stage, several fatty acid oxidation products are significantly affected in ASD. When only up-regulated genes were considered, reporter analysis identified alpha-fucosyl transferases and also metabolites in the phosphatidylinositol phosphate pathway as significantly affected. In the case of only down-regulated genes, metabolites in fatty acid oxidation, inositol phosphate metabolism, and nucleotide interconversion were identified.

At the MN stage, keratan sulfate degradation products were significantly identified as reporter metabolites. When considering only upregulated genes, the top 20 reporter metabolites consisted mostly of amino acids including glutamine, ornithine, lysine, isoleucine, and valine. Figures 3 and 4 show the metabolic set enrichment analysis and the Escher illustration for these metabolites, respectively. Escher illustrations of other significant metabolic pathways associated with these reporter metabolites is presented in Supplementary Figure S1.

**Figure 5.**
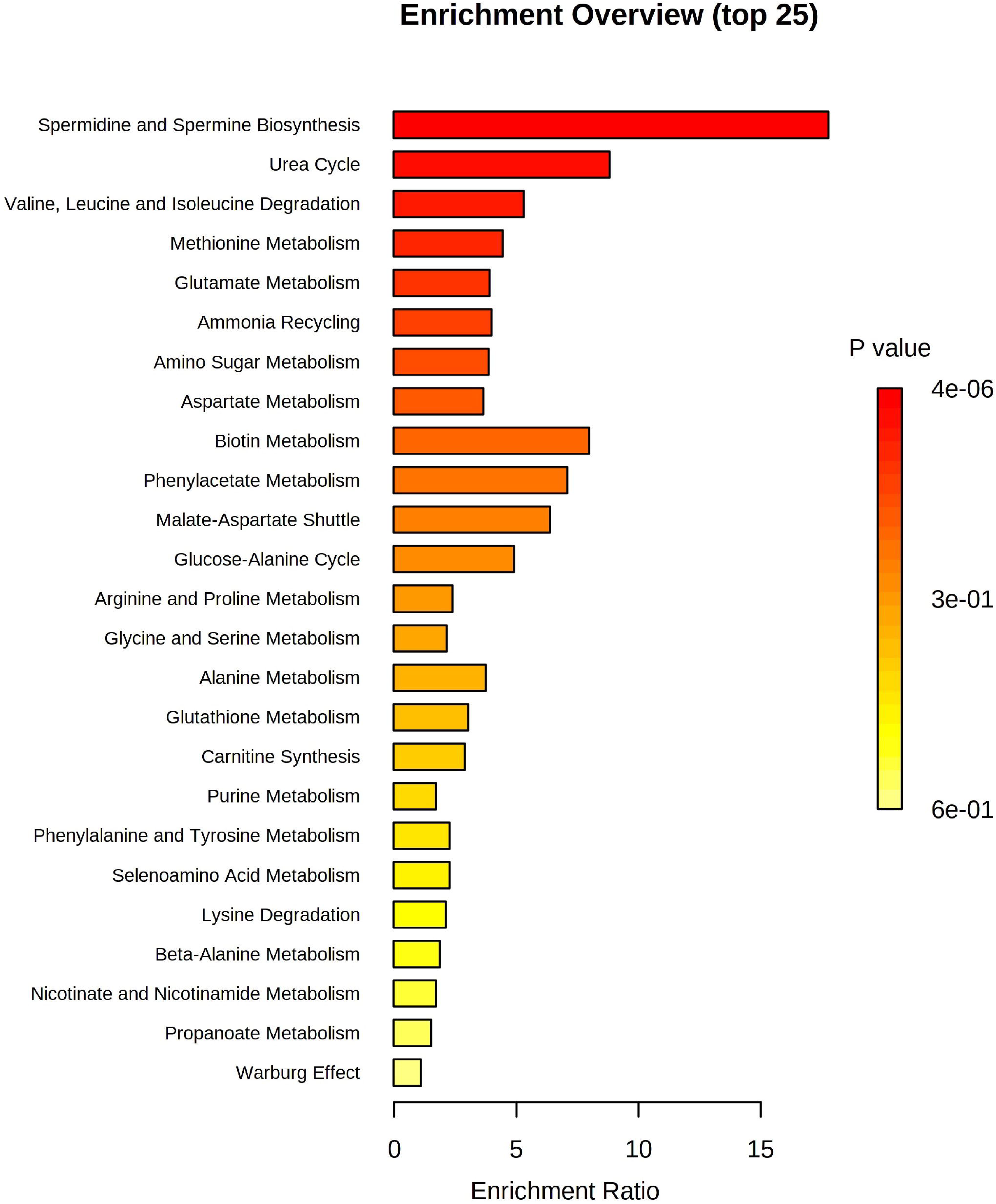
Reporter metabolite analysis on the mature neuron metabolic model. The results of metabolite set enrichment analysis of the top 20 reporter metabolites associated with the up-regulated genes are presented.

**Figure 6.**
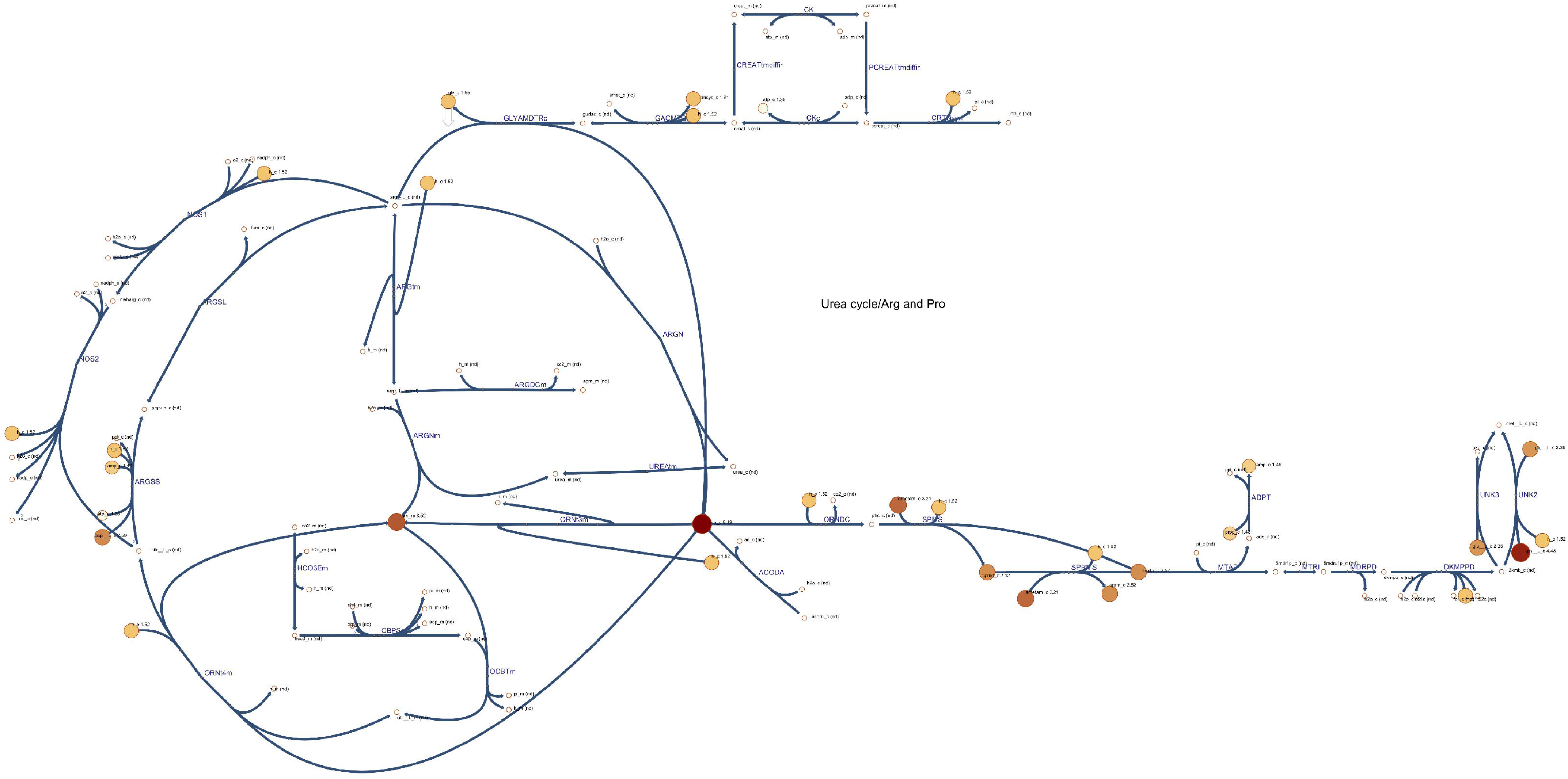
The Escher visualization of the results of the reporter metabolite analysis on the mature neuron metabolic model. Amino acids in the urea cycle are significantly altered between control and ASD models of the mature cortical neurons. Metabolites highlighted by the colored circles represent top reporter metabolites in the MN model when considering only upregulated genes. Numbers represent −log(*p*-value).

Reporter metabolites associated with only down-regulated genes mostly included lysosomal keratan sulfate degradation products. Notably, several keratan sulfate degradation products are significantly altered as predicted by reporter metabolite analysis.

**Figure 7.**
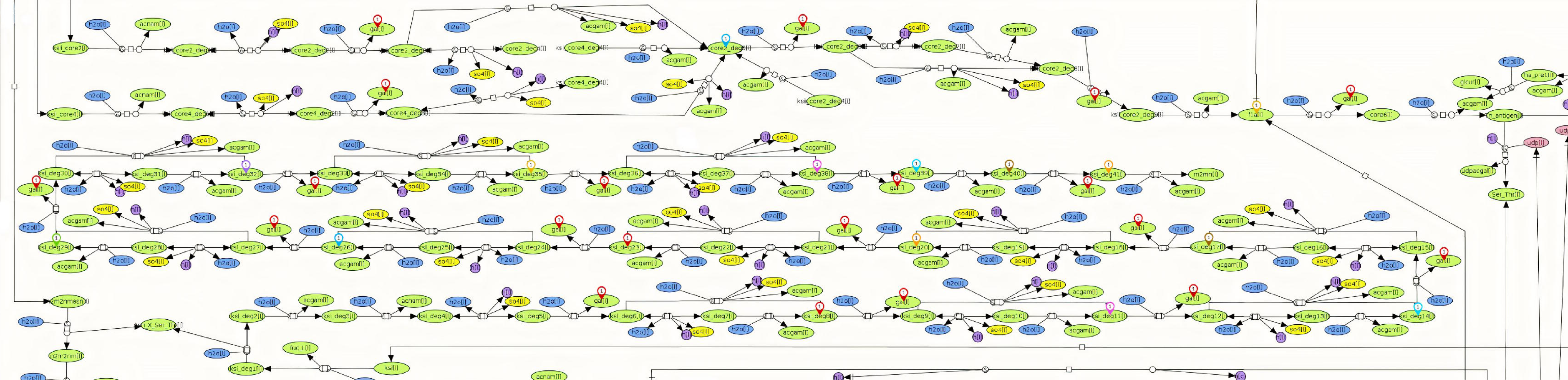
Lysosomal keratan sulfate degradation pathway. Multiple keratan sulfate degradation products are significantly altered in the mature neuron model as predicted by reporter metabolite analysis, indicated by location pins. This figure is taken from the Recon3D map provided on the Virtual Metabolic Human database (Brunk et al., 2018).

### Metabolic Transformation Algorithm and potential drug targets

Metabolic transformation algorithm sets to find those perturbations in the metabolic network that can globally shift the fluxes of a source state (*e.g.*, “disease”) in a way to recover a target state (*e.g.*, “healthy”). In this study, we used a recent implementation of MTA, namely rMTA, which was previously shown to have better performance (Valcárcel et al., 2019). Given a metabolic network reconstruction of the source state (here, ASD), the rMTA method requires two other inputs; differential gene expressions between the two metabolic states and a reference flux distribution (**v**^ref^) of the source state. The output of this algorithm is a list of reactions with their predicted robust transformation scores (rTS). Higher scores mean that the associated reactions are more favorable candidates in transforming the disease state into a healthy state, and are, therefore, potentially a more promising drug target for transforming the diseased state toward the healthy one. For each of the ASD models, 2000 sample fluxes were obtained and averaged to give the **v**^ref^. ASD metabolic models along with their respective differential gene expression analysis results and reference flux (**v**^ref^) were given to the algorithm and rMTA was run at the reaction level to obtain candidate reactions and their rTS values. Tables 3 and 4 show the top 10 reactions and their transformation scores at each developmental stage. Intriguingly, at the LNP stage, the most significant reactions predicted by rMTA are associated with phosphatidylethanolamine (PE) and phosphatidylserine (PS) metabolism. The top drug targets identified by rMTA at the mature neuron stage include phosphatidic acid phosphatase (PAP) and ATP: cytidine 5’ phosphotransferase. A list of reactions predicted by rMTA is available in Supplementary Data Sheet S6.

**Table 3.** Drug targets predicted by rMTA for neural progenitor model.

| Reaction Names | rTS score |
| --- | --- |
| Phosphatidylethanolamine Scramblase | 12.60 |
| Phosphatidylethanolamine Flippase | 12.60 |
| Phosphatidylethanolamine Transport | 12.54 |
| Phosphatidylethanolamine Flippase | 12.54 |
| Phosphatidylserine Flippase | 6.35 |
| Adenylate Kinase, Mitochondrial | 6.34 |
| Adenylate Kinase (GTP) | 6.34 |
| L-Lactate Dehydrogenase | 3.06 |
| Phosphatidylserine Transport by ABCa1 | 1.72 |
| (S)-Lactate:NAD<sup>+</sup> Oxidoreductase | 1.62 |
| Pyruvate Peroxisomal Transport via Proton Symport | 1.62 |
| Mitochondrial Carrier (Mc) TCDB:2.A.29.21.1 | 1.49 |
| Lactaldehyde Dehydrogenase | 1.31 |
| N-((R)-Pantothenoyl)-L-Cysteine Carboxy-Lyase | 1.24 |
| NADH Transporter, Peroxisome | 1.10 |
| Transport of NAD into Peroxisome | 1.10 |
| L-Aspartate Transport via Na, H Symport And K Antiport | 1.09 |

**Table 4.** Drug targets predicted by rMTA for mature neuron model.

| Reaction Names | rTS score |
| --- | --- |
| ATP: cytidine 5-Phosphotransferase | 21.5 |
| Phosphatidic Acid Phosphatase | 9.74 |
| Mitochondrial Carrier (Mc) Tcdb:2.A.29.21.1 | 9.09 |
| Saccharopine Dehydrogenase (NAD, L-Glutamate Forming), Mitochondrial | 8.79 |
| N6- (L-1, 3-Dicarboxypropyl)-L-Lysine:NAD<sup>+</sup> Oxidoreductase | 8.79 |
| Itaconate- Coenzyme A Ligase (GDP-Forming), Mitochondrial | 8.66 |
| Itaconate- Coenzyme A Ligase (ADP-Forming), Mitochondrial | 8.66 |
| Succinate-Semialdehyde:NADP<sup>+</sup> Oxidoreductase | 8.24 |
| Succinate- Coenzyme A Ligase (GDP-Forming) | 6.29 |
| ADP/ATP Transporter, Mitochondrial | 5.38 |
| Succinate- Coenzyme A Ligase (ADP-Forming) | 5.10 |
| Glutamate Dehydrogenase (NADP), Mitochondrial | 4.27 |
| Diacylglycerol Phosphate Kinase (Homo Sapiens) | 3.88 |
| Symport of Bicarbonate and Sodium | 3.69 |
| Succinate-Semialdehyde:NAD<sup>+</sup> Oxidoreductase | 3.58 |
| Glutathione Peroxidase | 1.67 |
| Malic Enzyme (NADP), Mitochondrial | 1.63 |
| Folate Reductase | 1.57 |

## Discussion

Autism Spectrum Disorders are complex neurodevelopmental disorders that are manifested through a variety of physiological and behavioral symptoms. Large cohort studies have resulted in numerous genetic mutations and transcriptional changes associated with ASD. However, any single mutation can explain only a small percentage of the cases. Nevertheless, there is a growing body of evidence that suggests ASD is associated with inborn errors of metabolism. In this study, we employed genome-scale metabolic network modeling to investigate metabolic dysfunctions in ASD. By leveraging available gene expression data, we reconstructed neuron-specific metabolic models for two developmental stages, namely, neural progenitors and mature neurons. The transcriptome data we used was obtained from a study in which the authors utilized a cortical neurogenesis protocol to differentiate IPSC-derived neurons. Cortical development has been previously implicated in ASD, with animal and post-mortem studies indicating macrocephaly, abnormal cortical organization, neuronal proliferation and migration in ASD brain. We therefore chose this dataset to further investigate the metabolic perturbations observed in neural progenitor and mature neurons in ASD. *in silico* gene knockouts allowed us to simulate the metabolic alterations in ASD in a systematic manner. Flux variability analysis revealed that in the ASD models at the neural progenitor stage, reactions associated with oxidative phosphorylation and protein N-glycosylation are significantly downregulated. Oxidative Phosphorylation is the major pathway for ATP production in neurons and several reports have previously indicated dysfunctions in OXPHOS and multiple complexes of mitochondrial electron transport chain in the blood, brain and muscles of ASD individuals (Anitha et al., 2012, 2013; Tang et al., 2013; Hollis et al., 2017). Among up-regulated reactions in our models, our results pointed to significant but less-studied pathways such as keratan sulfate (KS) metabolic processes, O-and N-glycosylation, and macroautophagy in ASD pathology. KS is a glucosaminoglycan and in humans, is most abundantly found in the connective tissue of brain and cornea. Several study lines suggest an important role for KS in the regulation of neural differentiation and development, synaptic plasticity, and axonal guidance. Enzymatic digestion of KS has been shown to promote the recovery of motor and sensory function after spinal cord injury in rats by enhancing axonal regeneration/sprouting. KS proteoglycans participate in forming perineuronal nets which are neuroprotective structures with antioxidant activity (Caterson and Melrose, 2018; Melrose, 2019). It is also reported that KS proteoglycans are upregulated following spinal cord injury (Imagama et al., 2011). Furthermore, previous studies have suggested an association between dysregulated macroautophagy and dendritic spinal density and pruning in postmortem ASD temporal lobe and also several mouse ASD models (Tang et al., 2014; Lieberman et al., 2020).

We also set to find the significantly altered metabolites that could act as potential biomarkers for ASD diagnosis. Keratan sulfate degradation products, intermediate products of fatty acid oxidation, and branched-chain amino acids consistently appeared in our analysis, which highlights the importance of these metabolites in ASD pathology. Lipid metabolism plays a major role in neuronal development and perturbations in fatty acid beta-oxidation and long chain fatty acid concentration have been frequently observed in ASD individuals (Clark-taylor and Clark-taylor, 2004; Tamiji and Crawford, 2010). Our results also show that there are alterations in spermidine and spermine, methionine, valine, leucine and isoleucine metabolism, and the urea cycle. Plasma and urine level alterations of these amino acids have been previously reported in several studies on ASD, however, their exact associations are yet to be described. Furthermore, mutations in the BCKD-kinase gene, which regulates branched-chain amino acid homeostasis, have been previously reported in ASD individuals (Zheng et al., 2017; Liu et al., 2019; Smith et al., 2019).

One of the interesting results of this study is the alterations in the spermidine and spermine biosynthesis in the ASD model of the mature neurons. spermidine and spermine are tightly regulated polyamines with extensive roles including chromatin structural modulation, neuroprotection, modulation of neuronal channels, and glia-neuron interactions. Polyamines regulate the excitatory synapses by blockade of glutamatergic receptors (AMPAR) and modulating Ca^2+^ flux and therefore spermine/spermidine imbalance can lead to neuronal excitotoxicity (Skatchkov et al., 2014). Mutation in the spermine synthase gene causes an X-linked intellectual disability syndrome (Snyder-Robinson Syndrome). Mutation in the *Drosophila* spermine synthase is also reported to cause retinal and synaptic degeneration. Spermidine catabolism produces reactive aldehydes and reactive oxygen species (ROS). The imbalance of these polyamines can therefore result in lysosomal dysfunction, impairment of autophagy, and oxidative stress that impairs mitochondrial function (Li et al., 2017).

Our results also predict perturbations in the Methionine metabolism. Methionine is one of the essential amino acids which is converted to S-adenosyl-methionine, S-adenosyl-homocysteine, and homocysteine, a pathway regulated by folate. S-adenosyl methionine is the main methyl group donor in the body and homocysteine is later converted to cysteine which is critical for glutathione synthesis. Therefore, the folate-methionine pathway plays a critical role in DNA methylation and synthesis, the maintenance of the redox balance, and the modulation of oxidative stress. Perturbations in the folate-methionine pathways have been previously reported in ASD in several studies. However, these studies are inconsistent and require further investigation to determine the exact association between this pathway and ASDs (Main et al., 2010).

Finally, rMTA suggested potential drug targets that may help recover the healthy metabolic state in ASD, namely phosphatidylserine (PS) and phosphatidylethanolamine (PE) transfer enzymes and phosphatidic acid phosphatase (PAP). Lower levels of PS have been observed in children with autism and reports suggest that supplements containing PS improve cognitive performance. PAP is a regulatory enzyme that plays a major role in lipid homeostasis, signaling, and synthesis of membrane phospholipids like PS and PE by catalyzing the dephosphorylation of phosphatidic acid (PA). PA is suggested to play an important role in membrane trafficking for neurite outgrowth and neurodevelopment (Ammar et al., 2014; Tanguy et al., 2019). Furthermore, Dysfunctions of PAP have been associated with neuropathy and neuroinflammation (Carman and Han, 2009; Carman, 2011). Other significant rMTA predictions include lactate dehydrogenase and oxidoreductase, pyruvate and NADH transport reactions, perturbations of which have been reported in ASD in several studies. Significant increase in lactate dehydrogenase and pyruvate levels in ASD led to Warburg phenomenon in these individuals (Khemakhem et al., 2017; Vallée and Vallée, 2018; El-Ansary et al., 2020). One interesting prediction by the rMTA is the ATP:Cytidine phosphotransferase. This reaction is regulated by two genes, uridine-cytidine kinase 1 and 2 (UCK1/UCK2). UCK1 and UCK2 phosphorylate uridine and cytidine to uridine monophosphate and cytidine monophosphate and are therefore crucial for the biosynthesis of pyrimidine nucleotides. A study has shown that plasma levels of uridine is significantly increased in autism, which is suggested to be a negative marker of methylation status (Adams et al., 2011). Studies on patients with CAD deficiency, which is characterized by developmental delay and intellectual disability, have shown that uridine supplementation leads to significant improvement in clinical symptoms (Rymen et al., 2020; Zhou et al., 2020). Another significant drug target predicted by rMTA is adenylate kinase (ADK). ADK is a key enzyme that synthesizes ATP and AMP from two ADP molecules (nucleotide interconversion) and is required for stable maintenance of cellular ATP and extracellular adenosine concentration, which in turn can affect several pathways involved in neuroinflammation, neuromodulation and synaptic transmission regulated by adenosine. As of now, there exists a report of rare mutations in the ADK gene associated with intellectual disability and autism (Najmabadi et al., 2011). Other noteworthy candidates predicted by rMTA include glutathione peroxidase, folate reductase, itaconate and succinate oxidoreductase, and coenzyme-A ligases.

## Conclusion

Autism spectrum disorders are a group of complex neurodevelopmental disorders with remarkable genetic heterogeneity and varying symptom severity. With the advent of genomic technologies and the availability of huge “omics” datasets, a systematic approach to studying complex disorders like ASD has proven increasingly useful. Research has shown that biomarkers of metabolic dysfunctions are consistently present in the blood, brain, muscle and urine of ASD individuals. Such dysfunctions include alterations in the mitochondrial electron chain complexes, oxidative phosphorylation and fatty acid oxidation and abnormal lactate/pyruvate ratios. In this study, by taking advantage of the available generic human metabolic model and transcriptome data, we reconstructed genome-scale metabolic network models of iPSC-derived cortical neurons at two developmental stages (neural progenitor and mature neuron). Following differential gene expression analysis, we simulated ASD metabolism by deleting significantly down-regulated genes between ASD and control in our neuron-specific model. We further aimed to study the metabolic alterations through several analyses including, FVA, GO term enrichment analysis, reporter metabolite analysis and rMTA. Our analyses confirm that oxidative phosphorylation and fatty acid oxidation are altered in ASD as reported in previous studies. Interestingly, our results also highlight the perturbation of two less-studied pathways in ASD, keratan sulfate degradation and O- and N-glycosylation. Finally, we propose several potential drug targets as predicted by our model to recover the healthy metabolism in ASD.

## Conflict of Interest

The authors declare that the research was conducted in the absence of any commercial or financial relationships that could be construed as a potential conflict of interest.

## Author Contributions

The original idea was presented by SAM. The development of the procedure and discussion of the results were done by MP and SAM. The manuscript was originally drafted by MP, and improved by SAM. The final version of the manuscript was read and approved by all authors.

## Funding

None declared.

## Acknowledgments

We would like to thank Mr. Meisam Yousefi for their kind help with the implementation of DESeq2 and CORDA2 algorithms.

## Supplementary Materials

**Supplementary File S1** Control metabolic network model of late neural progenitor cells.

**Supplementary File S2** Control metabolic network model of mature neurons.

**Supplementary Data Sheet S1** Exchange reactions and their boundaries.

**Supplementary Data Sheet S2** Differential gene expression analysis results.

**Supplementary Data Sheet S3** Reporter metabolite analysis results of neural progenitor stage.

**Supplementary Data Sheet S4** Reporter metabolite analysis results of mature neuron stage.

**Supplementary Data Sheet S5** FVA analysis. List of deleted genes, constrained reactions, fully decreased and increased reactions in ASD models of neural progenitor and mature neurons compared with control.

**Supplementary Data Sheet S6** List of reactions predicted by rMTA.

**Supplementary Figure S1** Escher illustrations of the metabolic pathways corresponding to the reporter metabolites associated with up-regulated genes at the mature neuron stage.

## Data Availability Statement

The context-specific metabolic network models are available as supplementary files. Publicly available datasets were used for model reconstruction in this study. These data were downloaded from Gene Expression Omnibus (GSE124308) and are available from: https://www.ncbi.nlm.nih.gov/bioproject/PRJNA511583

## Notes

https://www.ncbi.nlm.nih.gov/bioproject/PRJNA511583

### Competing Interest Statement

The authors have declared no competing interest.

