## Supplemental Figure for "Investigation of cell metabolism dysfunction in autism spectrum disorders using genome-scale metabolic network analysis of neuronal development"

Supplemetary Figure S1. Escher illustrations of the metabolic pathways corresponding to the reporter metabolites associated with up-regulated genes at the mature neuron stage. Reporter metabolites are enriched in Alanine, Methionine, Phenylalanine and Valine/Leucine/Isoleucin degradation pathway. Numbers indicate –log(p-value).

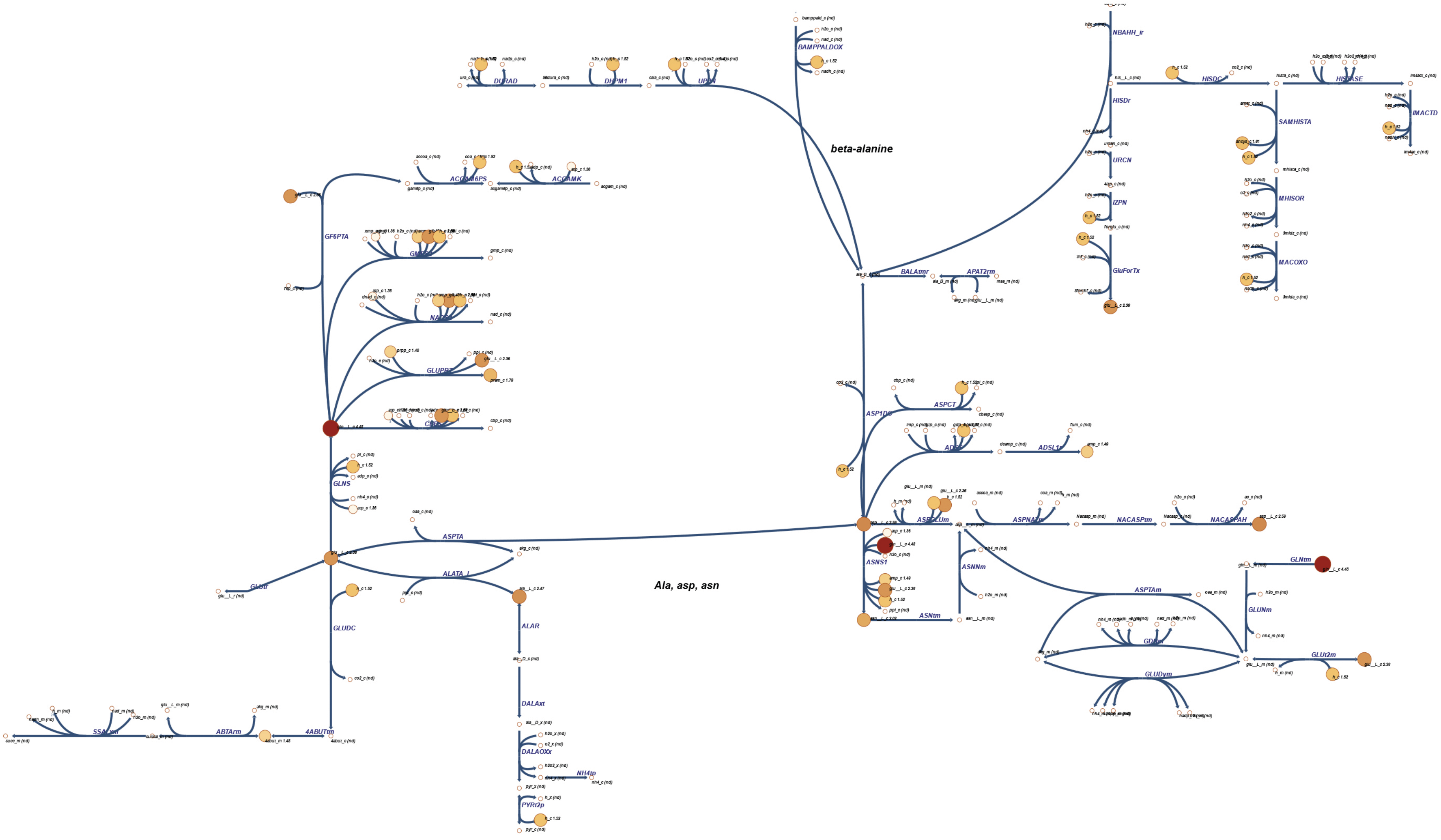

### *Methionine and cysteine metabolism*

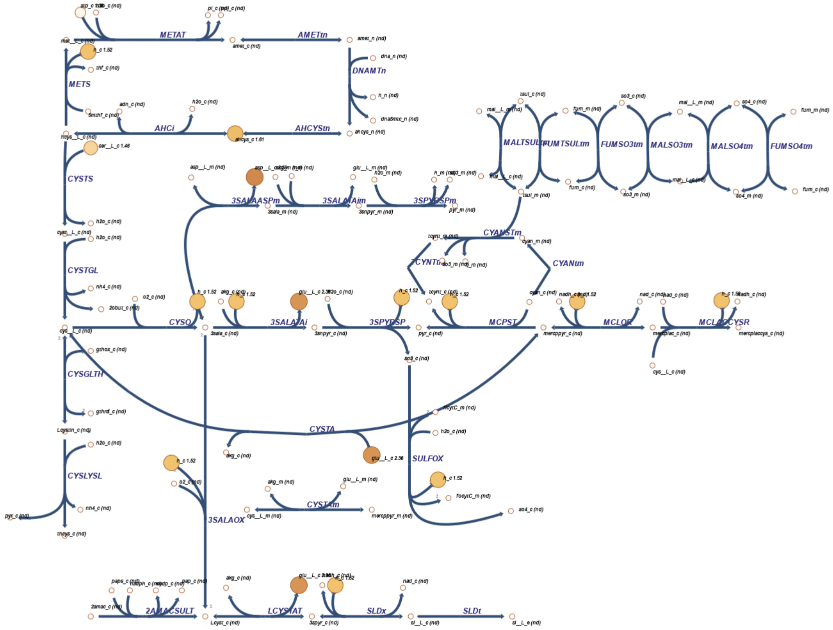

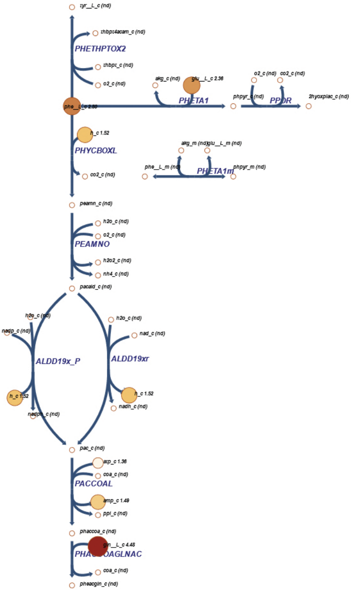
tyr__L_c (nd)
thbpt4acam_c (nd)
PHETHPTOX2
thbpt_c (nd)
o2_c (nd)
akg_c (nd)
glu__L_c 2.36
o2_c (nd)
co2_c (nd)
2hyoxplac_c (nd)
h_c 1.52
PHYCBOXL
co2_c (nd)
akg_m (nd)
glu__L_m (nd)
phe__L_m (nd)
PHETA1m
phpyr_m (nd)
peamn_c (nd)
h2o_c (nd)
o2_c (nd)
PEAMNO
h2o2_c (nd)
nh4_c (nd)
pacald_c (nd)
h2o_c (nd)
nadp_c (nd)
ALDD19x_P
h_c 1.52
nadph_c (nd)
h2o_c (nd)
nad_c (nd)
ALDD19xr
h_c 1.52
nadh_c (nd)
pac_c (nd)
atp_c 1.36
coa_c (nd)
PACCOAL
amp_c 1.49
ppi_c (nd)
phaccoa_c (nd)
gln__L_c 4.48
coa_c (nd)
pheacgln_c (nd)

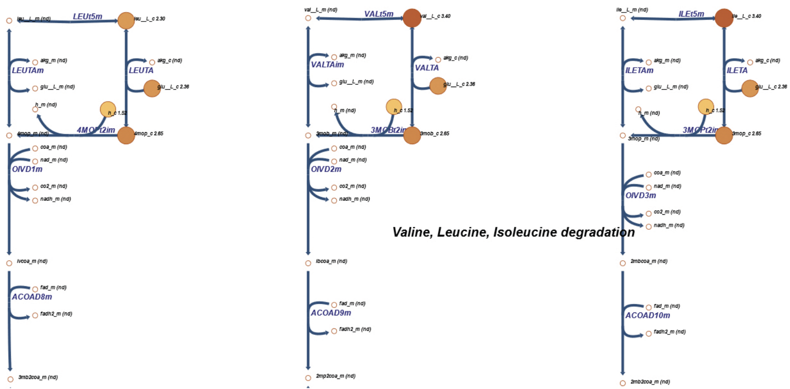
Valine, Leucine, Isoleucine degradation
LEUt5m
LEUTAm
LEUTA
4MOPt2im
OIVD1m
ACOAD8m
VALt5m
VALTAim
VALTA
3MOBt2im
OIVD2m
ACOAD9m
ILEt5m
ILETAm
ILETA
3MOPt2im
OIVD3m
ACOAD10m
